# Hippocampal ripples are distinguishable from aperiodic activity

**DOI:** 10.64898/2026.07.05.736595

**Authors:** James E. Kragel

## Abstract

High-frequency “ripple” oscillations support learning and memory across species, yet it has been argued that putative ripples in awake human recordings are false positives produced when algorithms misread aperiodic (1/*f*) fluctuations as ripple-band oscillations. We show that this conclusion arises from an artifact of evaluating detection algorithms on surrogate data containing only aperiodic activity. Ripple detectors are adaptive, setting their threshold from the amplitude statistics of the signal, so applying them to surrogate data that contains only aperiodic activity lowers the threshold and inflates false positives (median 62%). Adding real ripple-band events back to the surrogate corrects this threshold shift and eliminates most false detections across multiple standard algorithms. Using multi-variate classifiers, we show aperiodic fluctuations can reproduce the power of ripples but not their timing or spectral content. These findings indicate care needs to be taken when using surrogates to evalute ripple detection algorithms. Thus, under realistic signal properties, human hippocampal ripples remain distinguishable from aperiodic activity.

---

High-frequency “ripple” oscillations in the hippocampus support learning and memory in rodents, non-human primates, and humans (1–8). In humans, ripples are commonly identified with algorithms applied to both macro- and microwire recordings of the hippocampus obtained with intracranial electroencephalography (iEEG). A large body of work has related the timing of these events to memory (2, 5, 6, 8). If detectors commonly label other neural phenomena as ripples, then this prior work would need reinterpretation. van Schalkwijk and Helfrich (9) make such a claim. Using simulations, they suggest that putative ripples detected in awake human medial temporal lobe recordings are mostly false positives caused by detection algorithms misidentifying aperiodic (1/f) neural fluctuations as ripple-band oscillations. If valid, this concern would warrant reevaluation of the findings across species that make use of this approach.

In most studies, detection algorithms are not used to ask whether ripples exist but rather to mark when they occur so that their timing can be related to behavior or other neural events. Their use thus inherently assumes that the signals under consideration contain ripple events, in addition to other processes that generate activity in the ripple band. Ripples can readily be observed in hippocampal CA1, appearing as brief bursts (centered around 50 ms; 10) with a spectral peak in the ripple band, typically taken as 80–150 Hz in humans (11). Here we adopt an 80–120 Hz band, matching the detectors of van Schalkwijk and Helfrich (9). These properties matter because several kinds of activity overlap in the ripple band, including pathological high-frequency oscillations and interictal epileptiform discharges (IEDs) as well as ripples and gamma oscillations (11, 12). The question van Schalkwijk and Helfrich (9) raise is whether detectors pick out genuine ripples rather than one of these other sources, specifically aperiodic activity.

To ask whether aperiodic activity is misidentified by detectors, one needs a surrogate signal specifically designed to test this hypothesis. In their approach, van Schalkwijk and Helfrich (9) used aperiodic-only surrogates, assuming that false detections on these signals could be used to infer the performance of detection algorithms. However, in our view this surrogate is inappropriate to test this hypothesis, and it fails for two reasons.

First, ripple detectors are adaptive in nature. As commonly used, putative ripples are identified when power in the ripple band crosses 2 standard deviations (SDs) as determined by the entire signal distribution. By definition, this distribution includes a mixture of 1/f activity, ripples, and other events depending on the recording site. By using only 1/f activity as a surrogate, the mean and variance of the distribution decrease, which lowers the detection threshold (Fig. 1A). This more liberal criterion then counts a larger share of aperiodic fluctuations as ripples. The mean and variance of the surrogate distribution are thus shifted relative to the data the surrogate is meant to approximate, making inferences about detection performance invalid.

**Figure 1.**
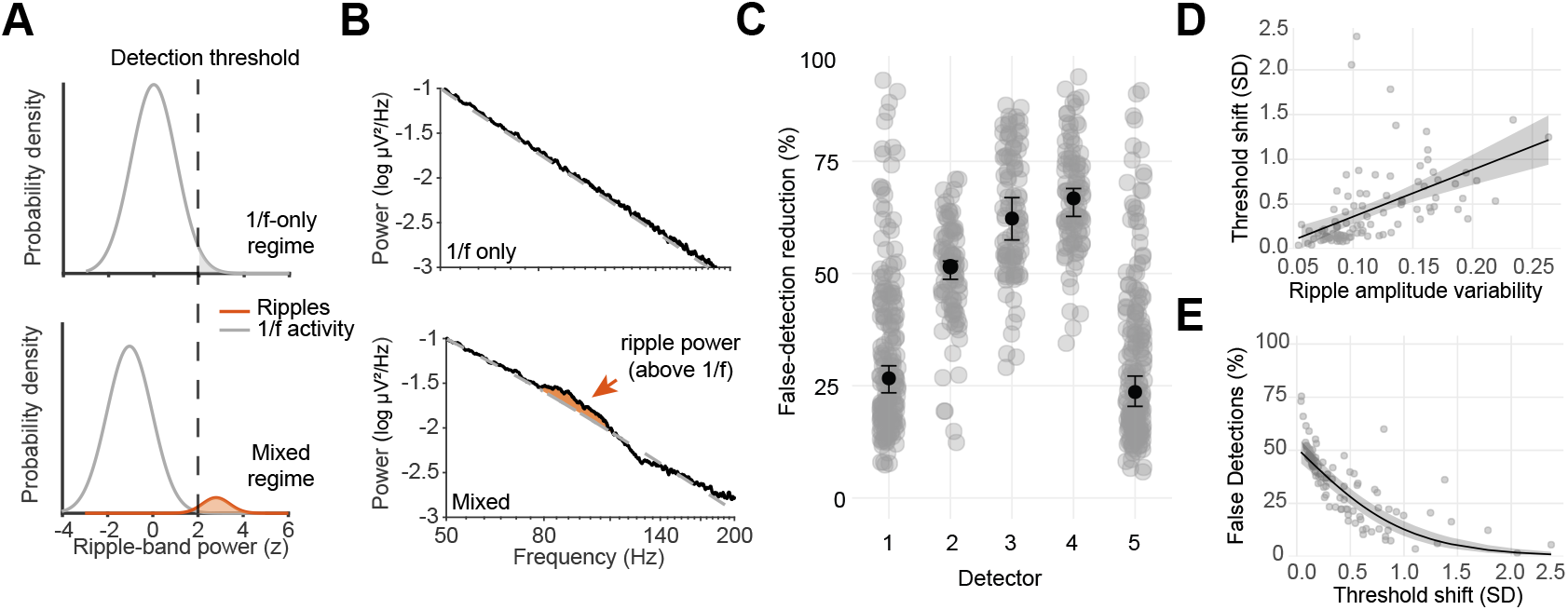
Excluding ripples from surrogate distributions inflates false detections. **(A)** Schematic illustrating ripple detection under 1/f-only (top) and mixed (bottom) regimes. In 1/f-only data, threshold crossings reflect aperiodic fluctuations, whereas in mixed data they reflect a combination of aperiodic fluctuations and ripples. **(B)** Representative hippocampal power spectrum (example contact from Study 1) showing localized ripple-band power above the fitted aperiodic (1/f) background in direct recordings (bottom) but not in matched aperiodic simulations (top). **(C)** Reduction in false detections when matched ripple-band events are restored to the surrogate, for each of the five detection algorithms. Points are individual channels (*N* = 99). Black markers show the median ±95% CIs. **(D)** Across channels, greater variability in ripple-band event amplitudes was associated with a larger shift in the detection threshold. **(E)** Larger threshold shifts removed a greater fraction of false detections. The fraction of surrogate detections surviving the inclusion of ripple-band events declined with the magnitude of the threshold shift. Points denote individual channels and shaded regions denote 95% CIs of predictions.

Second, detections in surrogate data cannot by themselves show that ripples observed in real data come from aperiodic activity. If 1/f activity were the source of putative ripples, the events detected in real and surrogate data should share the properties that define ripples such as peak frequency, duration, and their timing. This can be tested directly by comparing the events. If real and surrogate detections differ in these properties, the surrogate cannot account for real signals, which instead points to genuine high-frequency oscillations, ripples among them, as their source.

Here, we tested the viability of aperiodic-only surrogates in evaluating ripple-detection algorithms. To preview the findings, we first show that aperiodic-only surrogates inflate false detections, and that adding back ripple-band events removes most of them. We then use multivariate classifiers to distinguish real from surrogate ripple-band events, finding that the two primarily differ in duration and peak frequency.

To measure how much an aperiodic null inflates false detections, we applied all five detection algorithms used by van Schalkwijk and Helfrich (9) to hippocampal recordings. For each channel, we generated an aperiodic surrogate matched to that channel’s background and ran the same algorithms on it. The ripple-free surrogate produced a lower detection threshold and many more threshold crossings. We next added event waveforms from the hippocampal recordings back into the surrogate. These waveforms contain a mixture of ripples and other transients with ripple-band activity above the 1/f background, so the detectors now operated on signals resembling the ones they see in ordinary use.

Excluding ripples from the surrogate substantially inflated false detections. A median of 62% of detections flagged on the ripple-free surrogate went away once matched ripple events were added back (Figure 1C). This pattern held across all five detection algorithms, with the largest reductions for detectors relying solely on amplitude thresholding (Detectors 2-4) and smaller but consistent reductions for detectors with additional criteria such as event sharpness and duration (Detectors 1,5).

The number of false positives varied considerably across channels. Detection depends on high-amplitude events breaking above the background, so a channel whose ripple-band amplitudes span a wider range should be less prone to false positives. We tested this prediction for Detector 3, which uses ripple-band power and no other criterion. Channels with more variable ripple-band amplitudes showed greater threshold inflation (Fig. 1D, *β* = 2.81, 95% CI [0.90, 4.73], *χ*^2^(1) = 8.11, *p* = 0.004; within-subject *ρ* = 0.34). The increase in the SD used for thresholding accounted for this relationship. The effect of amplitude variability was no longer significant once SD inflation was included (*p* = 0.15), and SD inflation itself was strongly predictive of false detections (*p <* 0.001).

The larger the threshold shift produced by ripple-band events, the fewer surrogate detections survived (Figure 1E; *β* = −1.97, 95% CI [−2.37, −1.58], *χ*^2^(1) = 96.9, *p <* 0.001; within-subject *ρ* = −0.74). Thus, the aperiodic-only surrogate underestimates the threshold at which detectors operate in real recordings, where high-amplitude ripple-band events raise the amplitude distribution above the aperiodic background.

Having established that the aperiodic-only surrogate mischaracterizes the threshold at which detectors operate, we next turned to ask whether the surviving detections are oscillations. We therefore asked whether the detected events differ from the aperiodic excursions detected on a ripple-free surrogate. Applying the same detector to matched surrogates produced threshold crossings with clear ripple-band power (median 2.9 dB above the aperiodic background), close to that in real recordings (median 4.8 dB). The difference was not reliable across contacts (mean difference = 1.7 dB, 95% *CI* [0.00, 3.3], *p* = 0.07). Thus, fluctuations in 1/f activity can reproduce the observed ripple-band power during detected events.

Even if 1/f fluctuations match ripple-band power, they may not match the joint temporal and spectral structure of real events. For each detected event, we quantified seven waveform properties characterizing their frequency, duration, and power (see Methods). We then trained a logistic classifier to distinguish real from surrogate detections. It separated the two in held-out participants (AUC= 0.73, 95% CI [0.68, 0.77]; Fig. 2A). Duration and peak frequency carried most of the information for classification.Ripple power and other features carried little unique information (Fig. 2B). These results did not depend on a partic-ular detector. A classifier trained on one detector’s events generalized to others with only a small drop in performance (cross-detector AUC = 0.73, 95% CI [0.70, 0.76], versus 0.78, 95% CI [0.75, 0.81] within detector). Because real and surrogate events pass the same amplitude threshold, this separation comes from the temporal and spectral structure the aperiodic surrogate lacks, rather than ripple-band power.

**Figure 2.**
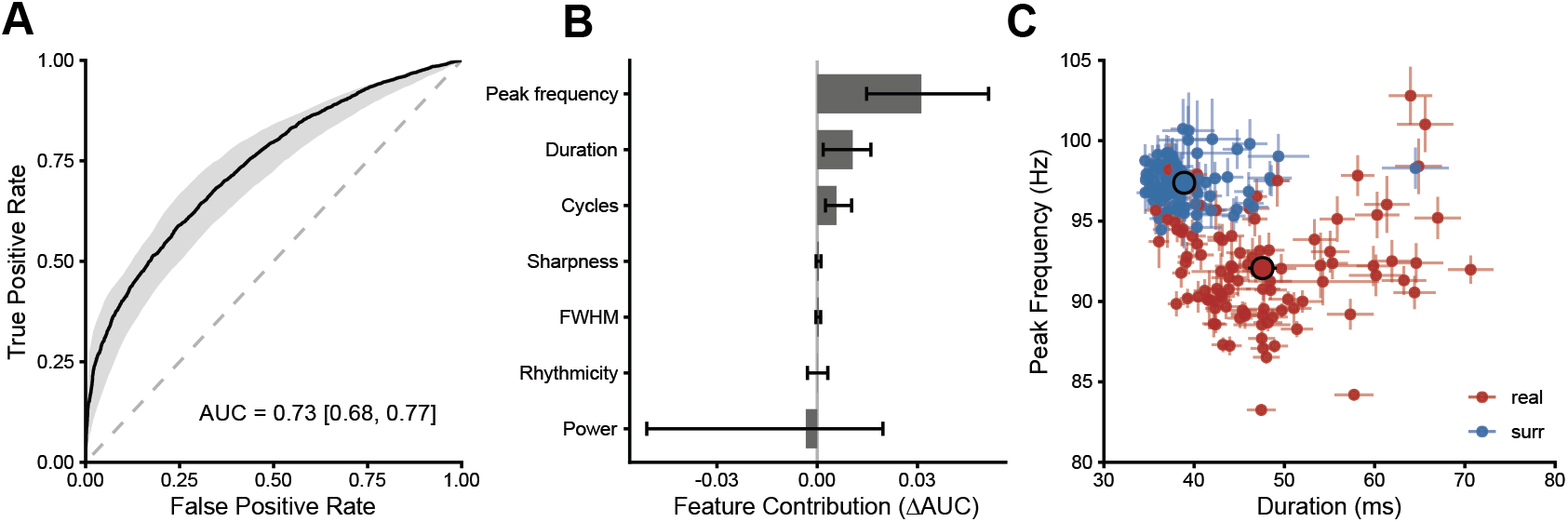
Detected ripple events are distinguishable from aperiodic surrogates. **(A)** Receiver operating characteristic curve for leave-one-subject-out cross-validation (detector 3). Shaded region shows the 95% CI (subject-level bootstrap). **(B)** Feature contribution, quantified as the drop in cross-subject AUC when each feature is removed. Error bars denote 95% CI. **(C)** Mean duration and peak frequency for real and surrogate events per channel (colored points). Black markers show group means, with lines denoting 95% CI.

What can a simulation like this tell us about whether a detected event is a real ripple? Only how a detector behaves on signals built from a particular set of assumptions. It gives no independent evidence that specific physiological events are present or absent. An aperiodic-only surrogate can show that a detector will cross a threshold when no oscillations exist, but not that the events it flags in real recordings share that origin. This limit applies to the work by van Schalkwijk and Helfrich (9) and to ours in equal measure.

Although the addition of ripples back into aperiodic-only surrogates leads to reduced false detections, it does not by itself disambiguate broadband transients from narrowband ripples. However, the comparison of spectral properties addresses this directly. Ripple-band events in hippocampal recordings, unlike those in aperiodic-only surrogates, carry the temporal and spectral structure that selection on amplitude alone cannot manufacture. Our findings therefore do not suggest that all detected events are necessarily putative ripples. Rather, they demonstrate that false-positive estimates derived from aperiodic-only simulations can overstate detector failures when the simulated signal distribution does not match the conditions under which detectors are applied to neural recordings.

In particular, because all recordings derive from epilepsy patients, a fraction of detected high-amplitude ripple-band events may reflect pathological activity rather than physiological ripples. We excluded events coincident with interictal discharges, and the 80–120 Hz detection band lies below the frequency range (*>*250 Hz) characteristic of pathological high-frequency oscillations (10, 12). Pathological events would, however, also raise the adaptive threshold and could contribute to the observed reduction. In the context of our findings, the spectral and temporal features that distinguished detected events from aperiodic excursions overlap with those shown to separate genuine hippocampal ripples from broadband interictal and pathological events (12), leaving their detailed physiological characterization to studies designed for that purpose.

Whether detected events reflect oscillations is a separate question, addressed by factors such as their spectral structure (e.g., Fig. 1B), phase autocorrelation, and waveform consistency. Our results indicate that whereas aperiodic surrogates can mimic changes in ripple-band power during detected events, real detections are distinguishable based on their joint temporal and spectral structure. Thus, the distinction between ripples and aperiodic activity rests on this structure rather than on power, the feature on which most ripple detectors rely.

These considerations point to what a surrogate must capture. Real recordings contain ripples mixed with aperiodic activity and other ripple-band events, and cortical amplitude fluctuations are heavy-tailed and non-Gaussian (13). A Gaussian, phase-randomized null omits this structure and understates the high-amplitude excursions a detector actually encounters, so detections on such a null cannot support inferences about detector behavior on real data. A surrogate is a tool for testing a specific hypothesis, and its construction determines which hypothesis it can test (14). Surrogates for assessing ripple detection must be validated for that purpose before false-positive estimates can be trusted. Our analysis is a small example of such validation rather than a complete generative model, and developing surrogates that reliably separate 1/f from oscillatory contributions remains an open goal.

The caution raised by van Schalkwijk and Helfrich (9), that task-related changes in high-frequency activity can reflect shifts in the aperiodic background, remains well founded and worth heeding when relating ripple activity to behavior. However, their analysis does not show ripples detected in real recordings are themselves aperiodic in origin. Under realistic signal statistics, hippocampal ripples remain distinguishable from the background against which they are detected.

## Methods

### Participants

#### Study 1

Data were drawn from previously published direct recordings of the hippocampus from epilepsy patients undergoing intracranial EEG while performing a scene recognition task (15). Data were collected at Northwestern Memorial Hospital (Chicago, IL) under protocols approved by the Institutional Review Board. Participants provided written informed consent. Study 1 contributes only the representative example in Figure 1B; all quantitative analyses use Study 2.

#### Study 2

Twelve patients (7 female, 45*±*16 years old [mean*±*SD]) undergoing presurgical monitoring for intractable epilepsy via stereoelectroencephalography were recruited to participate in this study. All participants had depth electrodes implanted in the hippocampus and additional structures to determine seizure onset zones. Data were collected at The University of Chicago Medical Center (Chicago, IL) and Northwestern Memorial Hospital (Chicago, IL). Research protocols were approved by the Institutional Review Board at both sites prior to data collection. Participants provided written informed consent to take part in the study. During the study, participants performed a visual field mapping task.

### Data acquisition and preprocessing

Intracranial EEG was acquired using a Nihon Kohden recording amplifier at 1–2 kHz with a scalp recording reference (Study 1) and with either a Nihon Kohden (at 1–2 kHz) or Neuralynx ATLAS (at 30 kHz) recording system, depending on the clinical site (Study 2). All recordings in Study 2 made use of a scalp reference. Recorded signals were bipolar re-referenced and downsampled to 2 kHz as needed. Preprocessing was conducted in MATLAB (MathWorks; Natick, MA) using custom scripts.

### Data-matched simulations of aperiodic activity

To evaluate detector performance under realistic conditions, we generated, for each recorded hippocampal channel, a surrogate matched to the aperiodic background and then incorporated ripple events observed on the raw recording. This procedure differs from a 1/f-only null in that the simulated signal contains both aperiodic activity and oscillatory events, including ripples, which is the regime in which detectors operate in practice.

#### Aperiodic background

For each channel, the power spectrum was estimated by multitaper spectral analysis `(pmtm,`time-bandwidth product *nw* = 5) in 30 s segments. Because hippocampal power spectra can exhibit changes in slope across frequencies, we fit an aperiodic model containing both a knee and an exponent to explain the observed spectra from 2–100 Hz. We used a robust, iteratively reweighted fit that masks oscillatory peaks before refitting the aperiodic components (16, 17). Because the fitting range overlaps the ripple band, the procedure relies on this peak masking to avoid absorbing oscillatory power into the aperiodic estimate. Any residual oscillatory power retained in the surrogate would, if anything, raise the detection threshold and make the reported reduction in false detections conservative. A surrogate reproducing the aperiodic background was constructed by shaping white noise in the frequency domain to match the fitted power spectral density. This process directly matches the aperiodic background across the ripple band while excluding any oscillatory peaks. Surrogates were constructed for each 30 s segment to match nonstationarity in the aperiodic background.

#### Restoring ripple-band events

To recover the amplitude distribution a detector would encounter in practice, observed event waveforms were extracted from detected events and added back into the surrogate at the empirical event rate, with a minimum inter-event interval of 200 ms. The amplitude of each event was scaled to match the distribution of detected event amplitudes for that contact. Because these were waveforms observed in actual recordings, the restored events comprise the mixture of ripples and other ripple-band transients present in the recording.

#### Contrast with prior work

The simulations of van Schalkwijk and Helfrich (9) correspond to the oscillation-free case. We applied each detector to both the original surrogate (without ripple-band events) and to the same surrogate with added events. Because the detection threshold is set from each signal’s own amplitude statistics, removing events lowers variability in the ripple-band signal and therefore the detection threshold. Correspondingly, adding ripple-band events back to the surrogate signals raises the threshold. Both analytic conditions used the same underlying surrogate, so they are matched in scale. Restored events were excluded from false-detection counts through temporal overlap.

#### Performance metrics

For each channel we used Monte Carlo sampling of surrogate data. The number of iterations was increased until reported measures changed less than 1% with increasing iterations. The reduction in false detections was computed as the proportion of detections in the original surrogate that were removed once the detection threshold was modified by ripple-band events. We additionally quantified the shift in the detection threshold in units of the standard deviation of the original surrogate.

### Ripple detection

We used the five detection algorithms described by van Schalkwijk and Helfrich (9). The following describes the primary detector (Detector 3), with the others differing in their filtering and threshold construction as specified in prior work. Putative ripples were identified with an amplitude-threshold detector applied to the 80–120 Hz band. The continuous signal was band-pass filtered to 80–120 Hz with a zero-phase IIR filter and the analytic amplitude was obtained from the Hilbert transform. The amplitude envelope was low-pass filtered at 10 Hz, edge samples were discarded, and the envelope was z-scored relative to its own mean and standard deviation. Candidate events were identified as peaks in the z-scored envelope exceeding 2 SD, with a peak width between 25 and 200 ms and a minimum separation of 500 ms between successive peaks. Detected events were further characterized by their peak and trough structure and a sharp-transient mea-sure, and events whose troughs fell within *±*1 s of a detected IED were excluded. Channels with excessive IEDs were excluded prior to analysis, leaving 99 contacts across 12 participants. For each retained event, the temporal extent used in the overlap-based performance analysis was defined by the surrounding samples at which the z-scored envelope last fell below the 2 SD boundary. Because the threshold is defined relative to each signal’s own amplitude distribution, the absolute amplitude corresponding to 2 SD is higher on signals that contain ripples than on aperiodic-only signals of matched slope; this is the mechanism by which the same nominal criterion produces fewer false detections when true oscillations are present.

### Interictal epileptiform discharge detection

IED detection was conducted based on established protocols (18). Preprocessing included band-pass filtering between 25 and 80 Hz, extraction of the signal envelope using the absolute Hilbert transform, and z-scoring. Events were detected when the envelope exceeded a threshold of 2 SD for a duration of 20–100 ms, and were terminated when the signal envelope did not exceed 2 SD. IEDs in close temporal proximity (1 s) were concatenated.

### Spectral analyses

Power spectral density (PSD) estimates used multitaper spectral methods. The aperiodic background was modeled with a model containing a knee and exponent, as described above. Ripple-band excess (Figure 1B) was quantified as the residual power in the 80–120 Hz band after subtracting the fitted aperiodic component, so that a positive residual indicates oscillatory power above the 1/f background rather than a mere increase in broadband high-frequency activity.

### Comparison of ripple-band events

To test whether ripple events detected in hippocampal recordings differed from those detected in aperiodic-only surrogates, we applied the primary detection algorithm to both timeseries and compared properties of the detected events. For each detected event, we estimated the multitaper power spectrum in a 200 ms window and computed two quantities relative to the aperiodic background fit for the corresponding 30 s segment. Ripple-band excess was the mean log power in the 80–120 Hz band above the fitted background. Spectral peak sharpness was computed from a higher-resolution spectral estimate (time–bandwidth product *nw* = 1.5) as the ripple-band peak frequency divided by the peak’s full width at half maximum, where the half-maximum was taken 3 dB below the peak and the crossing frequencies were found by linear interpolation. Peak sharpness is a scale-invariant measure equivalent to the quality factor (*Q*) of the peak, with higher values indicating a narrower, more resonant peak. Real and surrogate values were compared per contact with linear mixed-effects models as described below.

### Classification of detected events

To test whether events observed in real data differed from aperiodic surrogates, we compared events by applying the same detector to both types of data. For every detected event we computed seven features from a 200 ms window centered on the event trough: duration and cycle count (from the threshold-crossing extent and zero-crossings of the 80– 120 Hz signal); peak frequency, spectral peak sharpness, and the spectral peak full width at half maximum (FWHM; from the multitaper spectrum after subtracting the fitted aperiodic background); rhythmicity (defined by autocorrelation in average trough-locked responses); and ripple-band power above the aperiodic background. Features were *z*-scored using the mean and standard deviation of training data, computed within participant and applied to test data prior to classification.

We trained a logistic regression to classify events as real or surrogate and evaluated it with leave-one-subject-out cross-validation. The model was fit on all but one participant and used to predict held out events, iterating over participants. Discrimination was summarized by the area under the receiver operating characteristic curve (AUC), pooled across held-out predictions. The contribution of each feature was quantified as the decrease in cross-validated AUC when that feature was removed from the model, which reflects each feature’s unique contribution and is robust to collinearity among features. Confidence intervals were obtained by resampling participants with replacement (1000 bootstraps), so that uncertainty reflects the number of participants rather than the number of events.

To assess whether the distinction depended on a particular detection algorithm, we repeated the analysis for each of the five detectors and tested cross-detector transfer. A classifier trained on the events of one detector was applied, without refitting, to the events of every other detector, with features standardized using the training detector’s statistics. Classifier generalization was summarized among the detectors that separated real from surrogate events within detector, and its confidence interval was obtained by the same participant-level bootstrap.

### Statistical modeling

Analyses were performed in R. The reduction in false detections was summarized as the median across contacts, with 95% confidence intervals from bootstrap resampling. To test whether the magnitude of false detections depended on channel-level ripple properties, we fitted linear mixed-effects models with random intercepts per participant to account for repeated sampling of contacts within participants. Within-participant and between-participant components of each predictor were mean centered prior to modeling. For the channel-level relationship in Figure 1E, we modeled the fraction of surrogate detections remaining after threshold adjustment as a function of the change in threshold using a beta mixed-effects model `(glmmTMB;` 19) with a logit link and random intercepts per participant. Effects were evaluated by likelihood-ratio tests against nested models. The variability of event amplitudes and its mediation by envelope SD inflation were assessed in the same mixed-effects framework, separating within- and between-subject components by group-mean centering.

## Acknowledgments

The author thanks Joel Voss, Joshua Jacobs, and Philip Kragel for their comments on a previous version of this manuscript.

